# Novel mechanism of plasmid-DNA transfer mediated by heterologous cell fusion in syntrophic coculture of *Clostridium* organisms

**DOI:** 10.1101/2021.12.15.472834

**Authors:** Kamil Charubin, Gwendolyn J. Gregory, Eleftherios Terry Papoutsakis

**Affiliations:** Department of Chemical and Biomolecular Engineering & the Delaware Biotechnology Institute, University of Delaware, 590 Avenue 1743, Newark, DE 19713, USA

**Keywords:** *Clostridium ljungdahlii*, *Clostridium acetobutylicum*, syntrophy, heterologous cell fusion, hybrid cells, DNA exchange, DNA integration, plasmid DNA, antibiotic resistance

## Abstract

The evolution of bacteria is driven by random genetic mutations and horizontal gene transfer (HGT) of genetic material from other bacteria. HGT can occur via transformation, transduction, and conjugation. Here, we present a potential new mechanism of HGT which occurs in a syntrophic *Clostridium* coculture. We have previously shown that in syntrophic cocultures of *Clostridium acetobutylicum* and *Clostridium ljungdahlii*, the two organisms undergo heterologous cell fusion, which includes fusion of the peptidoglycan cell walls and membranes. Heterologous cell fusion facilitated a large-scale exchange of cytoplasmic protein and RNA between the two organisms, leading to the formation of hybrid bacterial cells containing cytoplasmic material of the two parent organisms. Here we present new evidence that cell fusion events also facilitate the exchange of plasmid DNA between the two organisms of the syntrophic coculture. Through the use of a selective subculturing process, we successfully isolated wild-type *C. acetobutylicum* clones which have acquired a portion of the plasmid DNA – containing the antibiotic resistance marker – from a recombinant strain of *C. ljungdahlii*. Fusion events led to formation of persistent aberrant hybrid cells with distinct morphogenetic characteristics. Furthermore, our data support the concept of a novel, interspecies, mechanism of acquiring antibiotic resistance. Since neither organism contains any known conjugation machinery or mechanism, these findings expand our understanding of multi-species microbiomes, their survival strategies, and evolution.

**IMPORTANCE:** Investigations of natural multispecies microbiomes and the field of synthetic and syntrophic microbial cocultures are attracting renewed interest based on their potential application in biotechnology, ecology, and medical fields. A variety of synthetic and natural cocultures have been examined in terms of their metabolic output, but relatively few systems have been interrogated at the cellular and molecular level. Previously, we have shown the syntrophic coculture of *C. acetobutylicum* and *C. ljungdahlii* undergoes heterologous cell-to-cell fusion, which facilitates the exchange of cytoplasmic protein and RNA between the two organisms, and leads to the formation of hybrid bacterial cells. Continuing this line of investigation, we now show that heterologous cell fusion between the two *Clostridium* organisms can also facilitate the exchange of DNA between the two organisms. By applying selective pressures to this coculture system, we isolated clones of wild-type *C. acetobutylicum* which acquired the erythromycin resistance (*erm*) gene from the *C. ljungdahlii* strain carrying a plasmid with the *erm* gene. Fusion led to persistent hybrid cells containing DNA from both parents but with distinct properties and morphologies. Moreover, we provide evidence for a novel mechanism of acquiring antibiotic resistance mediated by the syntrophic interactions of this system. This is a major finding that may shed light on a new mechanism of bacteria’s ability to acquire antibiotic resistance.

## INTRODUCTION

The evolution of bacteria and other single-cell organisms is facilitated through genetic mutations and horizontal gene transfer (1-4). In mutation-driven evolution, a cell acquires a random beneficial mutation(s), which is then passed on to daughter cells. In comparison, in the horizontal gene transfer (HGT) process, cells acquire beneficial mutations in the form of transferred DNA from other cells, often from a different species (1). The first evidence for HGT was provided in 1928, where pneumococci (in infected mice) exchanged virulence genes through the uptake of genetic DNA (1, 5). Since then, a great deal of gene transfer and genetic recombination examples have been demonstrated in the laboratory, while the genetic exchanges that occur in nature are believed to be widespread (1, 4). Horizontal gene transfer between bacteria can occur through various mechanisms, including transformation, transduction and conjugation.

Transformation is a process where competent cells take up genomic or plasmid DNA from their environment (1, 4). Natural competence has been observed under various environmental conditions, such as nutrient stress, involves 20 to 50 proteins (1), and has been documented in more than 80 species (3). Artificial competence can be developed in the laboratory through chemical treatment. The preparation of competent *E. coli* cells using CaCl_2_ treatment is an example (4). Transformation does not require cell-to-cell contact, as competent cells can take up foreign DNA from their environment. In transduction, DNA exchange between bacterial cells is carried out by a bacteriophage (4). This occurs when a portion of chromosomal DNA of a parent organism is accidentally packaged when a latent prophage excises from the genome. The phage containing the bacterial DNA can interact with and infect other cells, thus transferring the packaged DNA (4). Transduction does not require any cell-to-cell contact between cells either. Conjugation is the only known process where two bacterial cells have to interact physically in order to exchange DNA, which is mediated by a cell-to-cell junction through which the DNA passes (1). Conjugative DNA exchange has been characterized in most detail in Gram-negative bacteria, such as *E. coli*. The conjugative machinery is encoded on a conjugative plasmid or on integrative and conjugative elements (ICEs), a subset of which includes conjugative transposons (1, 2, 6). Conjugative plasmids have been identified in *Clostridium* species, an example of which includes the well-studied conjugative pCW3-like plasmids in *Clostridium perfringens* (7). Transfer of conjugative transposons from the Tn916/Tn1545 family into several *Clostridium* species from *E. coli* and *Enterococcus faecalis* has been reported, including *C. tetani, C. acetobutylicum*, and *C. beijerinckii*, with transfer between members of the genus also reported (8-10). It was reported that a Tn1545 self-mobilizing transposon, that conferred both tetracycline and erythromycin resistance, was transferred from *C. beijerinckii* to *Eubacterium cellulosolvens* (11). Importantly, no native conjugation systems have been found in either the autotrophic acetogen *C. ljungdahlii* (*Clj*) or the heterotrophic solventogen *C. acetobutylicum* (*Cac*) (2).

The syntrophic coculture system work we have recently reported (12) provides evidence for a new and never-previously described mechanism for HGT between bacteria. Plasmid DNA (p100ptaHalo) was successfully transferred from the *Clj*-ptaHalo strain – expressing the HaloTag protein (13) – to the wild-type (WT) *Cac* in a syntrophic coculture of the two organisms. Plasmid or other DNA transfer between these two organisms cannot take place through natural competency. First, there is no evidence that either *Cac* or *Clj* can become naturally competent (2, 3), and both are difficult to transform, largely through electroporation and at low frequencies. Second, as we describe here, the Restriction-Modification (RM) system of *Cac* prevents transformation with *Clj*-propagated plasmids. Since neither organism possesses an identifiable conjugation machinery, the DNA exchange we report here must have occurred through the newly identified heterologous cell-to-cell fusion in the coculture of the two organisms (12). We also provide evidence for a new form of interspecies-mediated *Cac* acquisition of antibiotic (erythromycin) resistance.

## RESULTS

### Enrichment and selection process for identification of *Cac* cells which have acquired plasmid DNA from the *Clj*-ptaHalo strain under coculture conditions

To determine if DNA can be transferred between *Clj* and *Cac* under coculture conditions, four biological replicates of parent cocultures of the *Clj*-ptaHalo strain, carrying the100ptaHalo plasmid, and the WT *Cac* were set up (Figure 1). Parent cocultures were set up in a non-selective coculture medium used previously (80 g/L glucose, 5 g/L fructose, no erythromycin) to encourage heterologous cell fusion (12, 14). p100ptaHalo carries the HaloTag gene and the erythromycin (Erm) resistance gene, *erm*, and can be propagated and expressed in both organisms (13). After 24 hrs, coculture samples were collected for subculturing and plating in selective media, chosen in order to enrich the cocultures in *Cac* cells that may have acquired the p100ptaHalo plasmid from the *Clj*-ptaHalo strain, while eliminating *Clj*-ptaHalo cells over time. The process involved subculturing in liquid medium for the first two passages, followed by plating on agar selection plates, and liquid subculturing of colonies from plates. Liquid subcultures were carried out in the selective Turbo CGM medium containing 80 g/L of glucose and no fructose, supplemented with 100 μg/mL of erythromycin. *Clj* cannot grow on glucose alone and its growth is in fact inhibited by such high glucose concentrations (14); fructose is the typical sugar on which *Clj* can grow. WT *Cac* cannot grow in the presence of 100 μg/mL of erythromycin, therefore erythromycin was used to eliminate WT *Cac* cells during the selection process. Liquid selection cultures were carried out in 100 mL bottles in an anaerobic chamber. Plating was done on 2xYTG plates, containing 5 g/L of glucose, no fructose, and 100 μg/mL of erythromycin.

**Figure 1.**
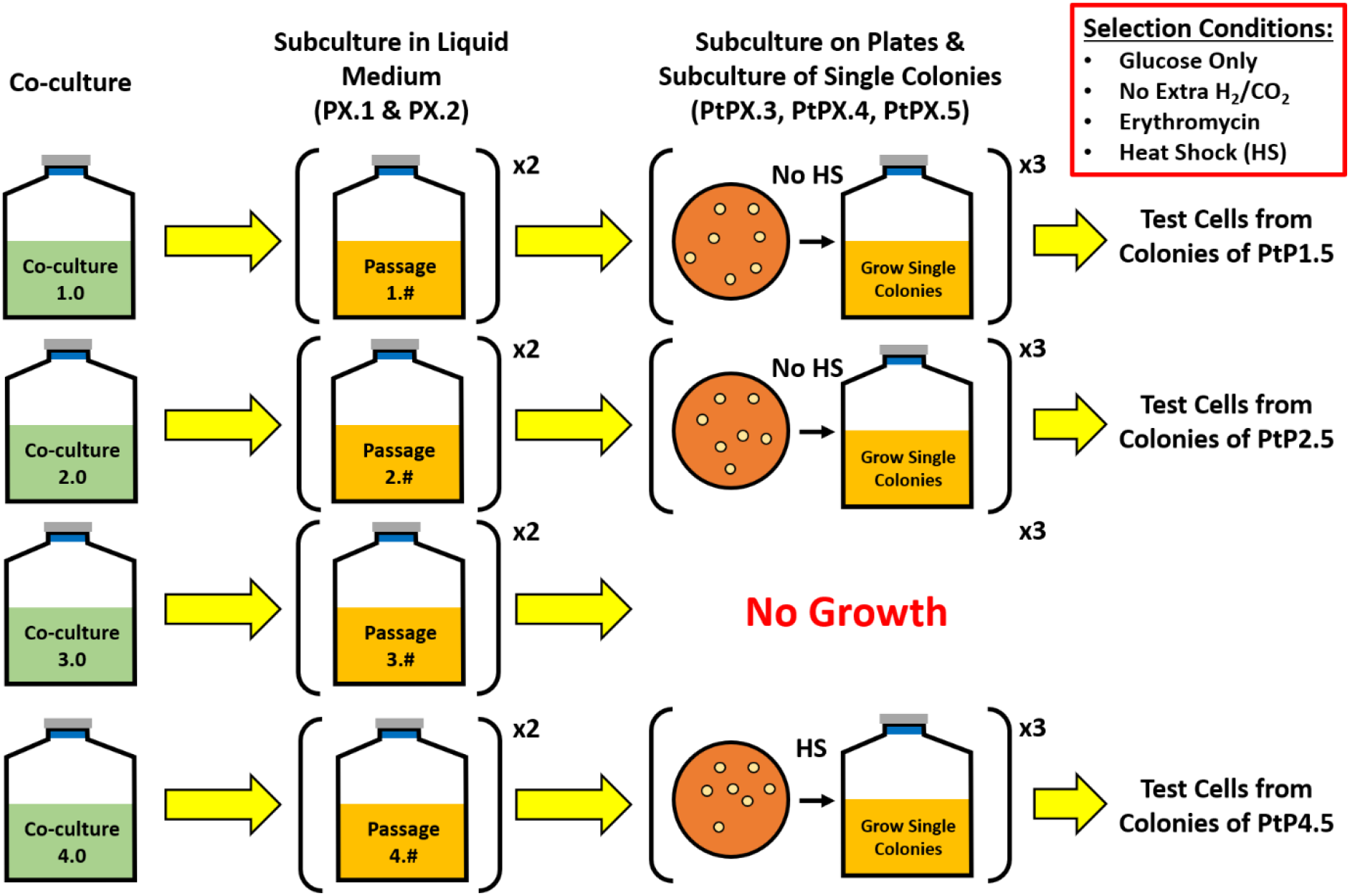
Summary of the selection procedure for isolating C*ac* strains, which have acquired p100ptaHalo plasmid DNA from *Clj*-ptaHalo cells in the coculture. Selection started with four parent cocultures (selection lines) on growth medium containing 80 g/L glucose and 5 g/L fructose but no erythromycin (Erm). Subculture passages 1 and 2 (PX.1, and PX.2; X represents the starting biological replicate of cocultures 1 through 4) were enrichment passages in selective liquid media containing glucose as the sole carbon source and Erm. Samples from the enriched cultures PX.2 were plated on selective plates (Erm, glucose) to identify and isolate single colonies of PtPX.3 strains. Selection subculture P3.2 did not survive on plates and was abandoned. Selection subcultures PtP1.3 and PtP2.3 (and all subsequent subcultures) could not survive heat-shock, indicating the lack of sporulation ability, and were subcultured without heat-shock for further analysis. Selective subculture PtP4.3 (and all subsequent subcultures) survived the heat-shock, indicating the presence of *Cac* cells capable of sporulation. Each subculture in liquid selection medium is represented as PX.#, where ‘X’ represents the parent coculture, while the ‘#’ represents each subsequent subculture (passage). Subcultures on selective plates are indicated as PtPX.#.

To start the selection process, the first selection subculture (PX.1; see methods and Figure 1 for numbering scheme) was cultivated in 25 mL of non-selective medium, containing 80 g/L glucose, 5 g/L fructose and no erythromycin for 24 hours. 5 mL samples from each PX.1 selection subculture were washed with Turbo CGM medium (80 g/L of glucose only, no fructose) and transferred to 25 mL (PX.2) of the selective liquid medium (glucose, Erm) to further enrich for plasmid-containing *Cac* cells. After 24 hrs, samples from subcultures PX.2 were streaked onto 2xYTG selection plates (plate subculture PtPX.3; glucose, Erm) to begin isolating and testing single colonies. The selection plates PtPX.3 developed colonies after 2 days of incubation at 37°C, except for selection plate PtP3.3 which did not develop any colonies (Figure 1). Thus, the P3.2 subculture was abandoned at that point. Eight to ten colonies were picked from each successful selection plate (PtP1.3, PtP2.3, & PtP4.3) and were subcultured in liquid selective medium. Half of the selected colonies were heat-shocked at 80°C for 10 minutes, per standard *Cac* culture practice, to select and grow *Cac* cells that have sporulated. Heat shocking kills *Clj* cells, as they are unable to sporulate. Colonies from plates PtP1.3 and PtP2.3 did not survive heat shocking, but grew in liquid selection medium if not heat shocked. Colonies from plate PtP4.3 survived the heat shock, and grew in the liquid selection medium. All colonies that grew in liquid selection medium were then streaked again on the 2xYTG selection plates (PtPX.4), and the process was repeated once more (passages PtPX.5), aiming to identify *Cac* cells resistant to erythromycin. Finally, four individual colonies from plates PtP1.5, PtP2.5, and PtP4.5 were grown in liquid selection medium, and analyzed using microscopy, flow cytometry for fluorescent protein (HaloTag) expression, metabolite analysis, and PCR assays to determine whether plasmid DNA was transferred from *Clj*-ptaHalo to the WT *Cac* cells during subculturing in selective media.

### Coculture mediated transfer of plasmid DNA from *Clj*-ptaHalo strain to WT *Cac*

The phenotype of subcultured cells and clones was assessed based on seven characteristics:

1. Survival of the heat-shock selection (*Cac* cells only)
2. Growth on 2xYTG plate surface (*Cac* cells only)
3. Growth on glucose only (*Cac* cells only)
4. Production of butanol & acetone (*Cac* cells only)
5. Growth with erythromycin (only possible if cells contain and express the *erm* gene)
6. HaloTag fluorescence (only possible if cells contain and express the HaloTag gene)
7. Production of isopropanol (coculture phenotype only (14): both *Cac* and *Clj* cells must be present or proteins from *Clj* present in *Cac* or *Cac* proteins present in *Clj*).

The phenotypic characteristics of the starting coculture partners (*Clj*-ptaHalo and WT *Cac*), the expected phenotype of *Cac* cells that carry the plasmid p100ptaHalo, and the phenotype displayed by the clones of the PtP4.5 plate are summarized in Table 1. Of the four parent cocultures (Figure 1), only individual colonies/clones from the PtP4.5 plate (4 originally and several more subsequently) showed all expected phenotypes of *Cac* cells that acquired p100ptaHalo from the *Clj*-*pta*Halo strain, with the exception of the plasmid-specific HaloTag fluorescence phenotype (Table 1). They: 1) survived the heat shock, 2) formed colonies on 2xYTG plate surface, 3) grew on glucose as a substrate, and 4) produced butanol and acetone (Figure S1), all of which are *Cac*-specific characteristics. Since these clones produce the characteristic solvents of *Cac* cells, the clones must contain these solventogenic genes, which are coded on the pSOL1 megaplasmid. Loss of pSOL1 results also in an asporogenous phenotype (15), and thus the ability of the clones to withstand the heat shock further confirms the presence of pSOL1. Significantly, no isopropanol was detected in cultures of this PtP4.5 strain despite the presence of 40 mM acetone (Figure S1), indicating the selection process worked at eliminating all *Clj*-ptaHalo cells from the original coculture. Any small amounts of acetone are readily converted to isopropanol by *Clj* cells (14). Erm resistance of the clones from the PtP4.5 plate can only be explained by the presence of the *erm* gene in the isolated clones. However, PtP4.5 clones did not show any red fluorescence when labeled with the red HaloTag ligand Janelia Fluor (Figure S2). Analysis of the earlier clones (from plates PtP4.3 and PtP4.4) showed that ∼30% of PtP4.3 cells exhibited red fluorescence, while PtP4.4 cells did not show any red fluorescence (Figures S2 and S3). Thus, during the 4^th^ selective passage (plate PtP4.4), cells lost the ability to produce sufficient levels of the functional HaloTag protein for fluorescence detection. To sum, these data show that at least a part of the p100ptaHalo plasmid, carrying the *erm* gene was transferred to *Cac*. The transfer of p100ptaHalo was examined in detail using PCR assays in the following section. To further confirm the identity of the clones from the PtP4.5 plate as *Cac* cells, we examined *Cac*-P4.5 clones morphologically using transmission electron microscopy (TEM). After 24 hrs of growth, *Cac*-P4.5 cells had the appearance of WT *Cac* cells (12): they displayed large translucent regions (granulose formed in preparation for spore formation) in their cytoplasm and a few fully formed *Cac*-P4.5 spores were observed (Figure 2). *Clj* cells (Figure S4) appear only as homogeneously electron dense (dark) vegetative cells (12), and no such cells were detected via TEM analysis.

**Table 1.** Phenotypic checklist of parent strains used for the starting coculture (*Clj-*ptaHalo and WT *Cac*), the expected phenotype of *Cac* cells which acquired the plasmid DNA (same as the *Cac*-ptaHalo strain), and the observed phenotype of isolated clones from plate PtP4.5

| *Characteristic* | *Clj*-ptaHalo | Wild-type *Cac* | *Cac*-ptaHalo (p100ptaHalo) | Clones from PtP4.5 Plate |  |
| --- | --- | --- | --- | --- | --- |
| *Heat shock survival* | No | Yes | Yes | Yes | ***Cac* Specific** |
| *Growth on glucose* | No | Yes | Yes | Yes |  |
| *Growth on 2xYTG plate surface* | No | Yes | Yes | Yes |  |
| *Production of butanol & acetone* | No | Yes | Yes | Yes |  |
| *Growth on erythromycin* | Yes | No | Yes | Yes | **Plasmid Specific** |
| *HaloTag fluorescence* | Yes | No | Yes | No |  |
| *Production of isopropanol* | No | No | No | No | **Co-culture Specific** |

**Figure 2.**
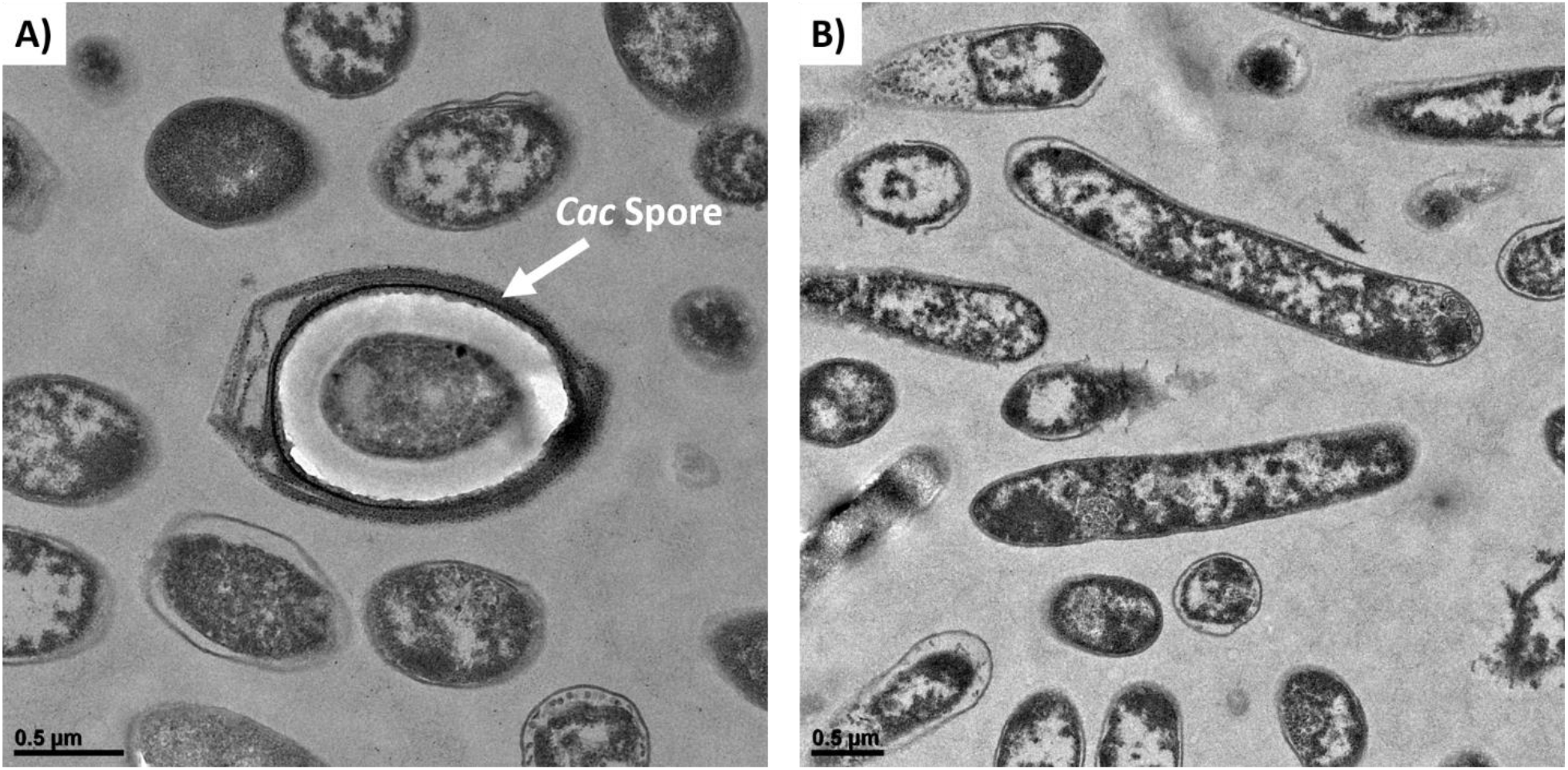
TEM imaging of clones from the PtP4.5 plate. (A) A fully formed *Cac* spore. (B) *Cac*-P4.5 cells showing the formation of granulose, which are the large white & translucent regions expected of sporulating wild-type *Cac* cells.

### PCR and PacBio analyses of the *Cac*-P4.5 cells identifies them as *Cac* cells containing both *erm* and HaloTag genes integrated into the *Cac* genome

First, *Cac*-P4.5 total DNA was tested for the presence of any *Clj*-ptaHalo genes. *Cac*-P4.5 cells were screened for five characteristic *Cac* genes (*adc, ctfA, ctfB, adhE1*, key solvent-formation genes encoded on the native pSOL1 megaplasmid, & *thl*, encoded on the main chromosome), and 3 *Clj* genes (*sadh, 23bdh*, & *rho*), as described (14). There is no DNA-sequence homology of the *Cac* genes in *Clj* and vice versa. Control PCR reactions with the pure WT *Cac* genomic DNA produced bands (∼100 bp) only with the *Cac*-specific primers (Figure 3A). Similarly, control reactions with the pure *Clj* genomic DNA produced bands (∼100 bp) only with the *Clj*-specific primers (Figure 3B). Thus, the primers designed for the assay showed very high specificity for the target organism. PCR-assay results from two colonies showed the presence of only *Cac* genes in the samples, with no *Clj* genes detected (Figure 3C).

**Figure 3.**
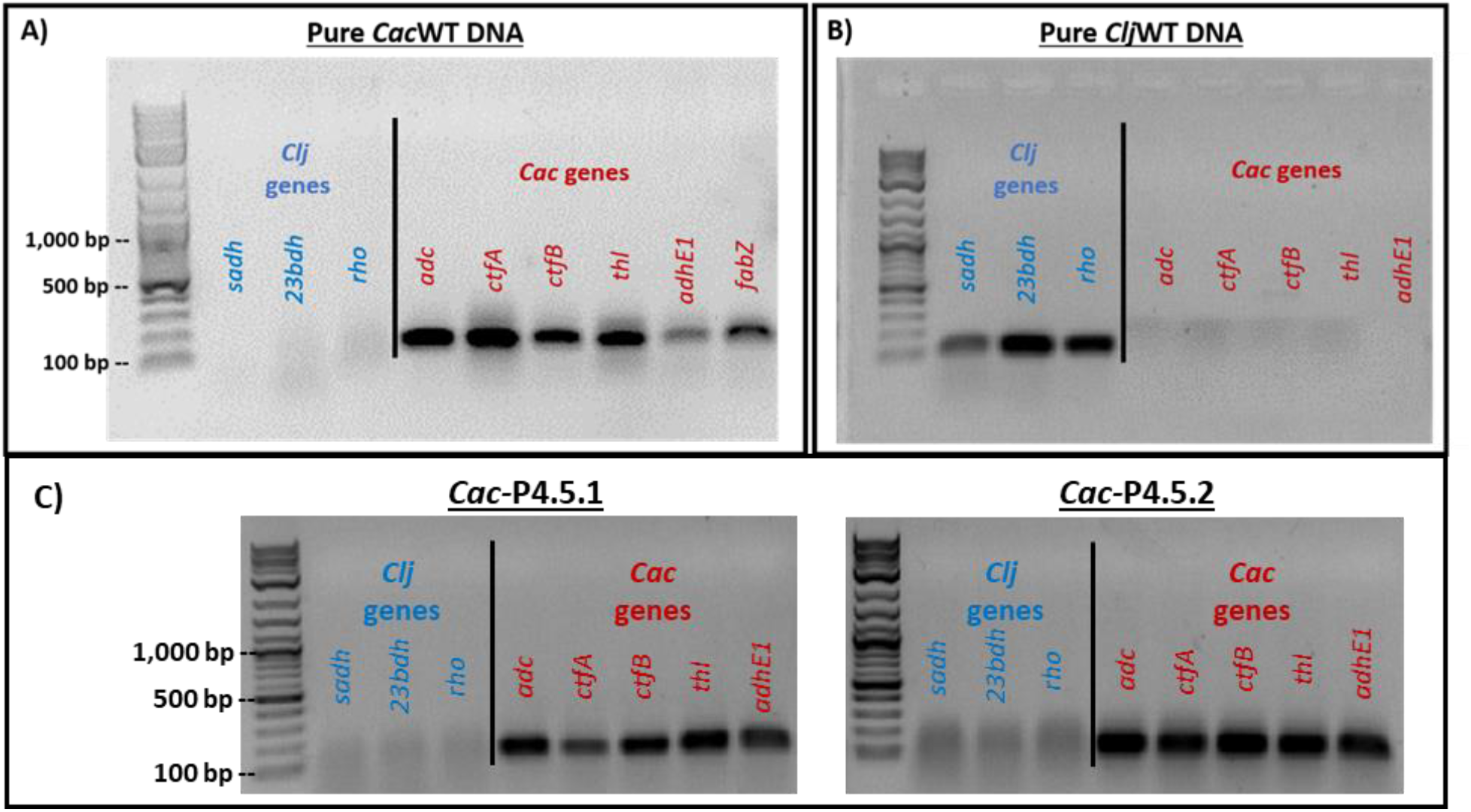
PCR control reactions with the selected primer sets on pure wild-type *Cac* and *Clj* genomic DNA. Three genes from *Clj* and five genes from *Cac* were targeted. (A) *Cac*-specific genes generated the expected ∼100 bp bands from the *Cac* genomic DNA template, while the *Clj-*specific primers did not. (B) *Clj-*specific genes generated the expected ∼100 bp bands from the *Clj* genomic DNA template, while the *Cac* primers did not produce any bands. Therefore, all primer sets chosen for each organism were highly specific for their target organism. (C) PCR assays used to test genomic DNA of the *Cac*-P4.5 clones for three *Clj*-specific and five *Cac-*specific genes. Results from two colonies (clones) are shown.

Three sets of primers were designed for the *erm* and the HaloTag genes to test the left (L) and right (R) flanks, as well as the middle (M) portions (Figure 4A). The flanking PCR reactions had one primer bind to either end of each gene, while the second primer would bind outside of the gene on the plasmid backbone (Figures 4A). Control PCR reactions for the *erm* gene showed identical results for both the total and p100ptaHalo DNA from the positive-control *Cac-*ptaHalo strain (Figure 4B). Total DNA from four individual colonies of the *Cac*-P4.5 clone was tested for the presence of the *erm* gene. The PCR reactions using the middle (M) and right (R) primer sets generated the expected bands (Figure 4C). The reaction using the left (L) set of primers generated a very faint band for three colonies, and no band from colony P4.5.3 (Figure 4C). Since the *Cac*-P4.5 strain was resistant to erythromycin, the *erm* gene sequence must be complete. Since the reaction using the left (L) set of primers did not generate the expected band, most likely the target sequence of the primer E1, which would bind to the left and outside of the *erm* gene, must be missing (Figure 4C). Since a faint band was visible for the (L) reaction, it is possible a few cells of the PtP4.5 clone might contain the full plasmid sequence, yet all efforts to isolate the p100ptaHalo plasmid from these clones failed. This implies that at least a portion of the p100ptaHalo plasmid, which contains the *erm* gene, was integrated into the *Cac*-P4.5 clone’s genome, or that a portion of the DNA was deleted from the p100ptaHalo plasmid. This hypothesis is further supported by the lower intensity of the M & R bands produced from *Cac*-P4.5’s DNA (Figure 4C), compared to bands generated from the *Cac*-ptaHalo’s DNA (positive control; Figure 4B). In both cases, the same amount of DNA template was used for the PCR reaction. *Cac*-ptaHalo cells contain multiple copies of the plasmid, and thus multiple copies of the *erm* gene, which resulted in the stronger PCR bands. If a portion of the p100ptaHalo plasmid integrated into *Cac*-P4.5’s genome, each cell of that strain would only carry a single copy of the *erm* gene, resulting in less PCR product, and thus bands with lower intensity.

**Figure 4.**
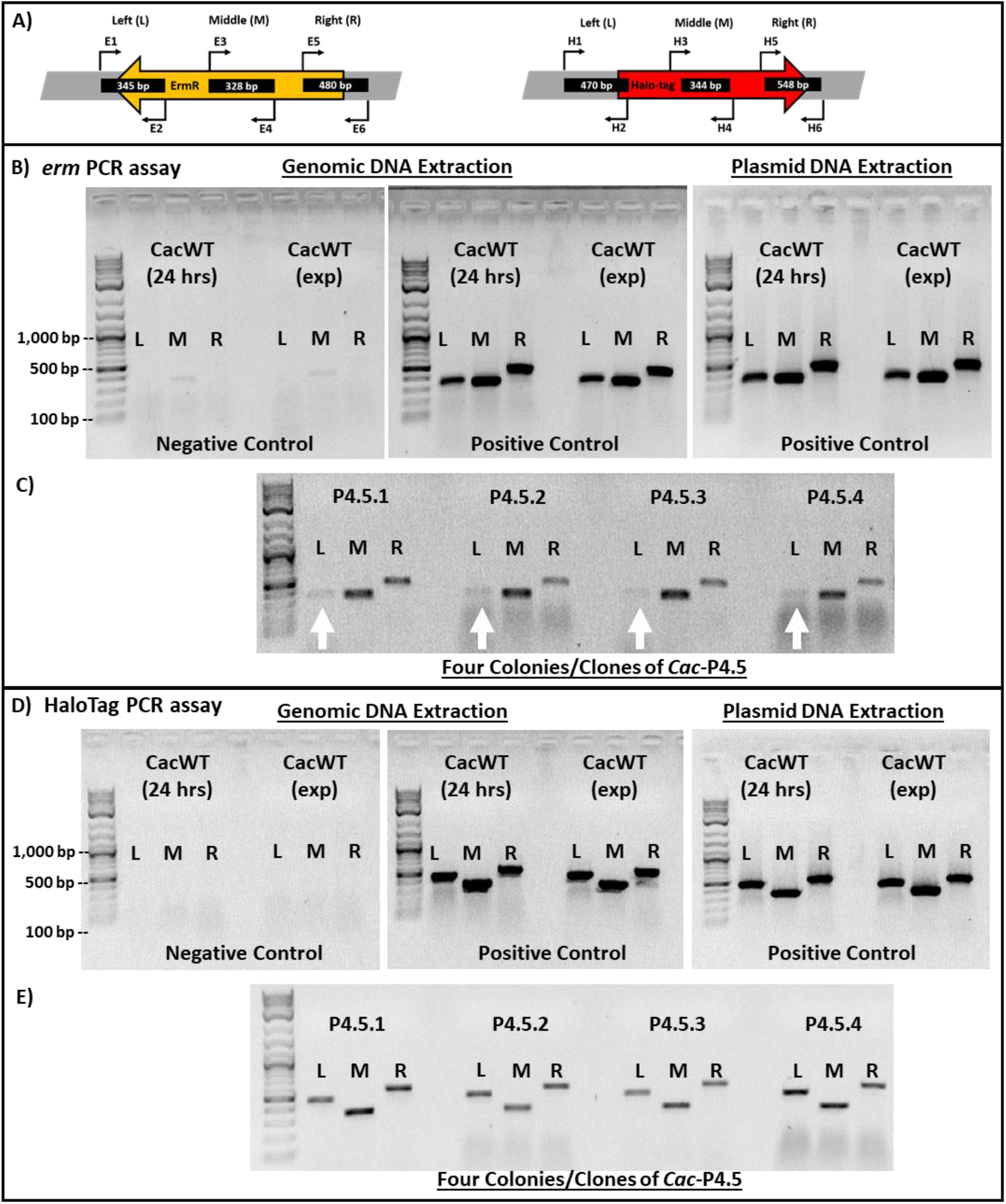
(A) Three primer pairs were designed to bind to the left (L), in the middle (M), and to the right (R) of the *erm* and HaloTag genes Primers E1 and H1 bind outside of each gene to the left, while primers E6 and H6 bind outside of each gene to the right, to test whether the original plasmid backbone was still present in the tested cells. (B) PCR assays to test the presence of the *erm* gene on the p100ptaHalo plasmid using negative control genomic DNA from the wild-type *Cac* or positive control DNA from *Cac*-ptaHalo. (C) PCR assays used to detect the *erm* gene using DNA from cultures initiated from four *Cac-*P4.5 clones (colonies). DNA from the *Cac*-P4.5 clone did not generate sharp, strong L bands (white arrows). The M and R reactions worked as expected. (D) PCR assays to test the presence of the HaloTag gene on the p100ptaHalo plasmid using negative control genomic DNA from the wild-type *Cac* or positive control DNA from *Cac*-ptaHalo. (E) PCR assays on the DNA isolated from the *Cac-*P4.5 clones produced all three expected bands, implying the entire gene is present. The bands in all four samples had a lower intensity compared to the positive control. The same amount of the DNA template was used in all *erm* and Halo PCR assays. Each individual colony selected for the assay were grown in the liquid selection medium until they reached optical density of 1.0-2.0. Cells collected from each culture were used for genomic DNA extraction.

The same *Cac*-P4.5 DNA samples were tested for the presence of the HaloTag gene as well. Control PCR reactions for the HaloTag gene showed identical results for both the total and p100ptaHalo DNA from the positive-control *Cac-*ptaHalo strain (Figure 4D). All three PCR reactions (L, M, R) of the HaloTag gene assay produced the expected bands in all four DNA samples from the *Cac*-P4.5 strain (Figure 4E). Why were the *Cac*-P4.5 cells non-fluorescent (Figure S2) when labeled with the HaloTag ligand if they did carry the HaloTag gene? The bands generated from *Cac*-P4.5’s DNA had a lower intensity, compared again to the positive-control *Cac*-ptaHalo DNA sample (Figure 4D). As shown in the *erm* gene assay, this again implies that the *Cac*-P4.5 clone contains fewer HaloTag gene copies compared to the *Cac*-ptaHalo strain. This provides additional evidence for plasmid DNA integration into the *Cac*-P4.5 clone’s genome. A single copy of the HaloTag gene would most likely not be able to produce enough of the HaloTag protein per cell to reach the detection threshold by flow cytometry. Examining the flow cytometric (Figure S2) and microscopic data (Figure S3) of PtP4.3 and PtP4.4 cells, it appears that there is a gradual loss of the free plasmid which is apparently complete by the 4^th^ selection passage (plate PtP4.4).

Analysis of PacBio sequencing data from a P4.5 clone identified six reads that contained the *erm* gene or the HaloTag gene, three of which contained two full copies of the p100ptaHalo plasmid, with no homology to the *Cac* genome. A full copy of the plasmid was identified inserted in the chromosome at position 2,734,720, with ∼3000 bp and ∼6000 bp on either end of the read aligning to the genome, while two and a half copies of the plasmid were inserted into the chromosome at position 671,974, with ∼7000 bp of the read corresponding to the *C. acetobutylicum* genome. While the low number of reads identified containing plasmid DNA was surprising, it is consistent with PacBio analyses of lineages P1.5 and P2.5, discussed below, which have higher numbers of plasmid reads and from which intact plasmid was isolated. Analysis of sequenced reads from *Cac*-P4.5 suggests that the plasmid is integrated into the chromosome at two loci. The integration of the plasmid into the genome, along with a small number of reads containing only plasmid DNA, supports the PCR analysis data, as well as the flow cytometric and microscopy data. The PacBio data suggests that there are very few copies of the *erm* and HaloTag genes present in each cell. Therefore, bands of lower intensity would be generated after PCR due to fewer copies of the genes as compared to the control (Figure 4). Additionally, very low levels of the HaloTag protein would be present which explains the lack of signal in the presence of the HaloTag ligand in both the flow cytometric and microscopy data (Figures S2 & S3). The PacBio data were also used to successfully assemble, *de novo*, the complete *Cac* genome including the pSOL1 megaplasmid, thus supporting the phenotypic characteristics of the *Cac*-P4.5 cells. No *Clj* DNA could be identified in the sequence data of the *Cac*-P4.5 cells. To support these findings, we also sequenced clones from P4.3 and P4.4. The P4.3 clone had six reads that contained at least 500 bp of the p100ptaHalo plasmid, with at least one end of the read corresponding to *Cac* genomic sequences (insertions at 525,552; 1,325,197; 1,823,332; 2,702,390; 2,779,170; and 3,413,786). Four reads from P4.4 contained at least 500 bp of the p100ptaHalo plasmid with at least one end corresponding to the *Cac* genome (insertions at 349,946; 780,069; 980,388; and 2,204,734). Furthermore, in contrast to P4.5, there were a few hundred reads containing *Clj* DNA (but no reads with both *Clj* and p100ptaHalo plasmid DNA) in P4.3, but considerably fewer reads in P4.4. Taken together, these data suggest a dynamic integration – under the antibiotic selective pressure – of the plasmid into the *Cac* genome at each subculture passage, as the loci of integration events changed and the number of integration events decreased with each passage. The presence of *Clj* DNA in P4.3 and P4.4 suggests that at earlier stages of lineage 4 DNA from both organisms co-existed. This is pursued further below.

In summary, p100ptaHalo plasmid DNA originating from the *Clj*-ptaHalo strain successfully transferred into WT *Cac* cells during the coculture. Over the course of the selection process, the p100ptaHalo integrated into *Cac*’s genomic DNA, creating the *Cac*-P4.5 clone. These integration events as well as detection of portions of the p100ptaHalo plasmid containing the *erm* and Halotag genes in the PacBio sequencing data explain the observed phenotype of the *Cac*-P4.5 clone. We confirmed that the *Clj* genome does not contain native conjugation machinery or mobile genetic elements that could be responsible for plasmid transfer to *Cac* by searching the reference genome using online Mobile Element Finder and ICEFinder tools (16, 17). No mobile genetic elements were detected in the *Clj* genome. Importantly, the reference *Clj* genome does not contain any ICEs, indicating that the most likely cause of DNA exchange is via the observed cell fusion, and not other known means of DNA transfer such as conjugation.

### Clones PtP1.5 and PtP2.5 show evidence for the formation of persistent hybrid bacterial cells containing chromosomal DNA from both *C. acetobutylicum* and *C. ljungdahlii*

Subculturing of the other two cocultures resulted in the isolation of clones PtP1.5 and PtP2.5 which exhibited a more complex and unexpected phenotype compared to the phenotype of *Cac*-P4.5 clones described above. Single colonies of subculture plates PtP1.5 and PtP2.5 were cultured in the liquid selection medium for analysis. Their phenotype was not consistent with either pure *Clj* or pure *Cac* strains (Table S1). Starting with the 3^rd^ subculture, the PtP1.5 and PtP2.5 clones could not survive the heat-shock, as would be the case for *Clj*. The same two clones had *Cac-*like phenotypes, including growth on glucose only, butanol and acetone production, and colony growth on the 2xYTG plate surface (Table S1). Furthermore, both clones were Erm resistant, and HaloTag fluorescent. Most surprising was the production of high concentrations of isopropanol (∼80 mM titers) by the PtP1.5 clone (Figure S5A & B), which should only be possible in the coculture of *Cac* and *Clj* or if hybrid *Cac*/*Clj* cells persist (14). In the coculture, *Cac* and *Clj* cells were found to undergo heterologous cell-to-cell fusion, which facilitates the exchange of proteins and RNA (12). Based on these data, these clones should contain characteristic genes of both *Cac* and *Clj* cells.

Detailed PCR analysis was performed as above to test for the presence of *Clj* or *Cac* genes and the *erm* and HaloTag genes. In all three tested individual colonies from PtP1.5 plate, the PCR data show that these clones carry all three *Clj* genes and all five *Cac* genes, as well as the full *erm* and HaloTag genes (Figure 5) with PCR-product band intensities similar to those produced with genomic DNA from the *Cac*-ptaHalo strain (positive control). We note the results shown in Figure 5A would also be obtained from genomic DNA extracted from a mixture of *Cac* and *Clj* cells. These data then suggest that the PtP1.5 clones contained multiple copies of the *erm* and HaloTag genes, thus suggesting the presence of an intact p100ptaHalo plasmid. As shown in Figure S5C & D, PtP1.5 clones contain a small population of cells (1.3% to 8.0%) expressing the HaloTag protein at levels that result in a strong enough red signal to be detected by flow cytometry. Therefore, at least some of the cells were able to maintain and express the p100ptaHalo plasmid over the course of the selection process. To confirm this, we were able to easily isolate the full p100ptaHalo plasmid from both sets of PtP1.5 and PtP2.5 clones. If hybrid *Cac*/*Clj* cells were only transient, it would be expected that over the course of multiple cell divisions, proteins from the other organism would be diluted, degraded, eventually disappearing over time, and the hybrid cells would revert back to their original identity. PacBio sequencing analyses of these two clones supports the formation of persistent hybrid cells. Mapping of the reads to the *Clj* and *Cac* reference genomes via PacBio SMRT Link analysis demonstrated that the full chromosome of each species (including the *Cac* pSOL1 native megaplasmid that carries all the genes for solvent formation) was present in the clonal populations, as well as the full p100ptaHalo plasmid. In lineage P1.5, there were six reads with both p100ptaHalo DNA and *Cac* DNA detected. In lineage P2.5, there were 44 reads that contained both p100ptaHalo plasmid DNA and *Cac* DNA, and five reads that contained both p100ptaHalo DNA and *Clj* DNA. Additionally, 259 reads were identified with >500 bp of p100ptaHalo plasmid from P1.5 and 7,016 reads from P2.5, which is consistent with isolation of intact plasmid from each of these lineages. The higher number of plasmid reads identified in P1.5 and P2.5 also supports the low number of reads found in P4.5, from which intact plasmid could not be isolated.

**Figure 5.**
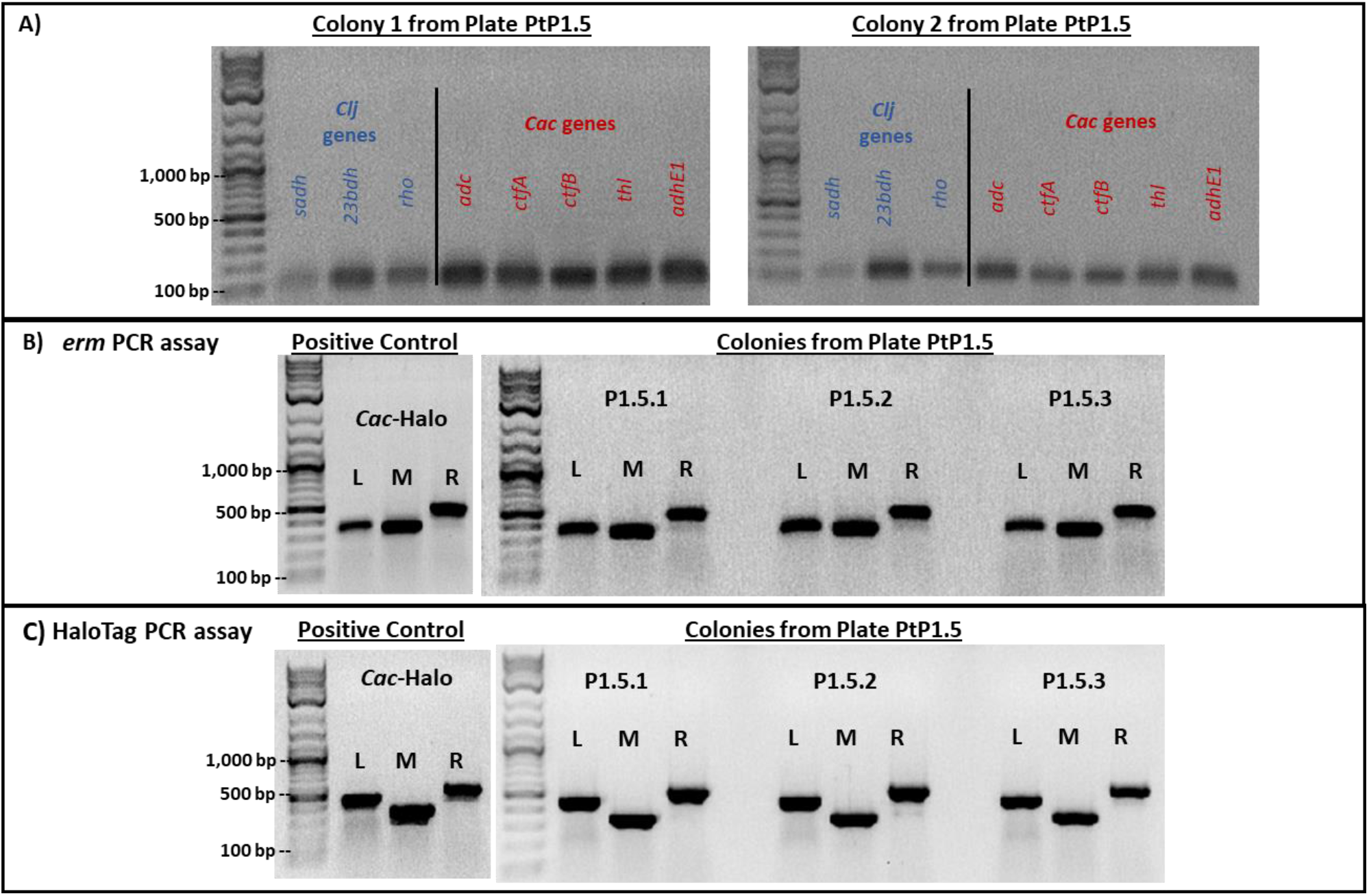
(A) PCR assays to test genomic DNA of PtP1.5 clones for three *Clj* (blue) and five *Cac* (red) genes. Results from two individual colonies are shown. Both were positive for the *Clj* and *Cac*-specific genes. (B) PCR assays used to detect the *erm* gene in PtP1.5 clone. DNA from the positive control produced the expected reaction products (bands), with strong intensity for the left (L), middle (M), and right (R) reactions. DNA extracts from three PtP1.5 colonies produced all three reaction products (bands); band intensities were similar to those of positive control. (C) PCR assays used to detect the HaloTag gene in PtP1.5 clones. DNA from the positive control produced the expected reaction products (bands), with strong intensity for the left (L), middle (M), and right (R) reactions. DNA from three PtP1.5 colonies produced all expected reaction products (bands). Band intensity was similar to that observed in the positive control. The same amount of DNA template was used in each PCR reaction. PtP1.5 colonies were picked from a PtP1.5 plate and subcultured in liquid selective medium until cells reached an optical density of 1.0-2.0. Harvested cells were used for DNA extraction.

### TEM examination of PtP1.5 and PtP2.5 clones identifies aberrant cell morphologies

To probe the nature of clones from plates PtP1.5 and PtP2.5 further, we examined them morphologically after 24 hrs of culture using TEM. PtP1.5 cells had an ambiguous morphology, that resembled neither pure *Cac* nor pure *Clj* (Figure 6A & B, Figure S4 for comparison). TEM imaging of PtP2.5 cells also showed ambiguous and unexpected morphologies (Figure 6C & D). While most of the examined PtP2.5 cells displayed the mixed *Cac-Clj* morphology of hybrid cells, a few cells had an enormous length of more than 10 μm. This morphological phenotype indicates that cell division does not function normally in cells containing protein, RNA, or DNA material originating from the two organisms. These TEM images of PtP1.5 and PtP2.5 cells further support the possibility that a chromosomal DNA exchange took place between *Clj*-ptaHalo and WT *Cac* cells in the coculture, or during the subculturing process. While both chromosomes are present, key native properties of both *Cac* and *Clj* cells were lost. If pure WT *Cac* cells persisted, they would not have been Erm resistant, they would have been able to sporulate and thus resist heat shock, and would thus have developed the advanced sporulation forms like those seen in Figure 2. If pure *Clj* cells had persisted, they would not have been able to grow on 2xYTG plate surfaces and would display the characteristic homogeneous electron dense vegetative form in TEM analysis (12). Yet, the primary metabolic capabilities of *Cac* cells were preserved, as well as the unique metabolic phenotype of heterologous cell fusion: ability to form isopropanol and 2,3-butanediol without acetone and acetoin accumulation in the medium (12, 14). These data further support the concept of persistent hybrid cells containing both chromosomes.

**Figure 6.**
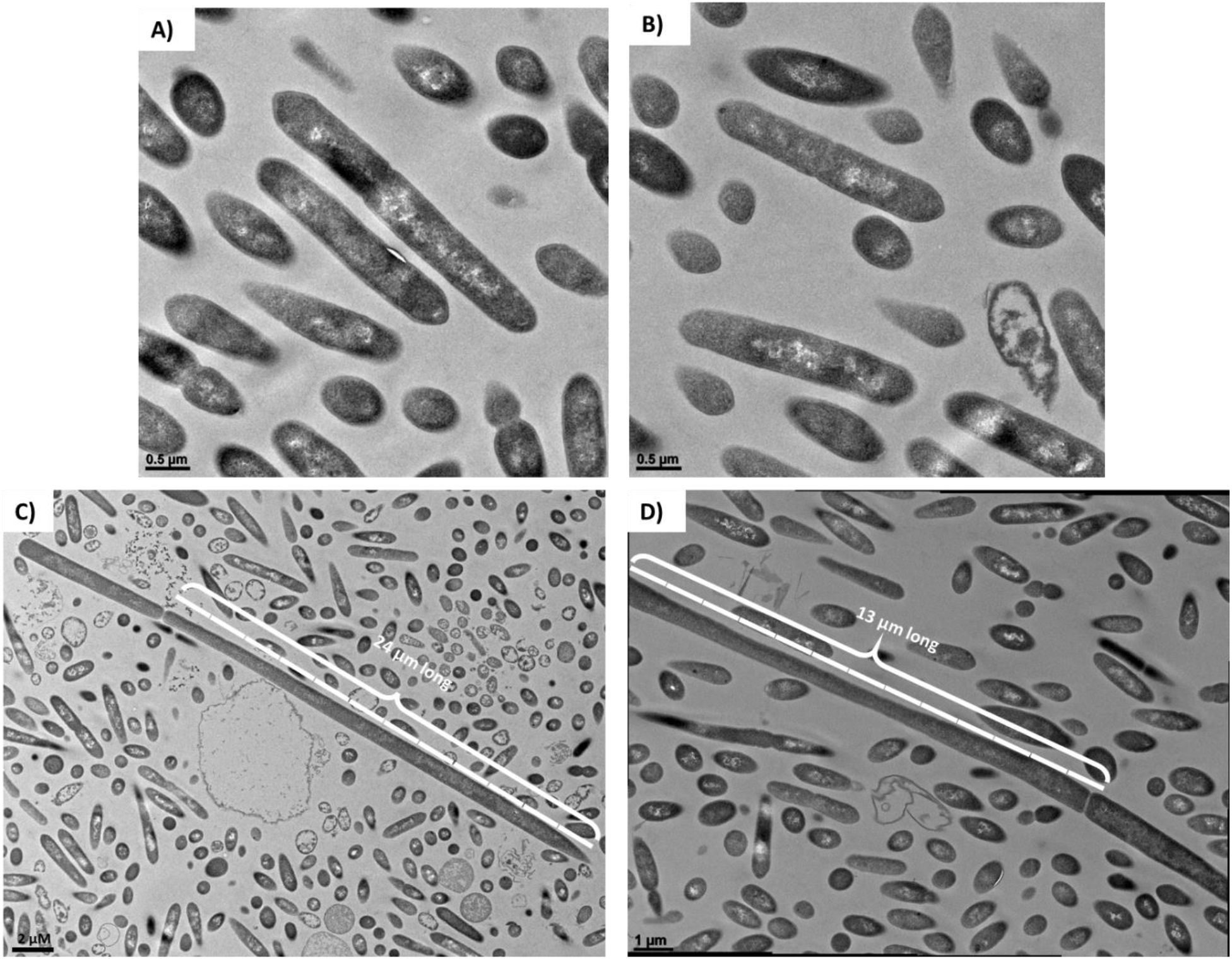
(A) and (B) TEM imaging of PtP1.5 cells after 24 hrs of a culture initiated from a single colony on a selective plate. All cells had an ambiguous morphology, displaying some differentiation and granulose formation (*Cac*-specific phenotype), but not as pronounced and defined as would be expected of pure *Cac* cells (compare to Figures 2 and S4). (C) and (D) TEM imaging of PtP2.5 cells after 24 hrs of a culture initiated from a single colony from a selective plate. All cells had an ambiguous morphology, displaying some differentiation and granulose-like formation similar to the PtP1.5 cells. A few cells in the sample had an enormous length of >10 μm, which is >5 times the size of a normal *Cac* or *Clj* cell (2 μm length).

The enlarged cells observed in the PtP2.5 clone (Figure 6C & D) were also observed in the PtP4.3 clone (Figure S3), but not in PtP4.5 (Figure 2). This is consistent with the PacBio sequencing data whereby a small number of reads aligning to *Clj* DNA were detected in P4.3, but considerably fewer reads (10-fold) were detected in P4.4, and no *Clj* DNA was detected in P4.5. An interpretation would be that expression of *Clj* genes coded on the detected *Clj* DNA in P4.3 would result in expression of *Clj* proteins that interfere with the morphological development of *Cac*.

## DISCUSSION

The goal of this study was to examine the possibility of DNA transfer between *Clj*-ptaHalo and WT *Cac* cells in co-culture (Figure 7), in addition to the large-scale exchange of protein and RNA reported previously (12). It was assumed that if any DNA exchange does occur, it would be a less frequent event, compared to the protein and RNA exchange. Thus, a selection process was designed to identify any WT *Cac* that acquired plasmid DNA from *Clj*-ptaHalo (Figure 1). The selection process generated cells with two unique phenotypes. First, the *Cac*-P4.5 clone was isolated with the expected phenotype of a *Cac* cell (heat-shock survival, butanol & acetone production, no isopropanol nor 2,3-butanediol production, and expected TEM morphology) carrying the *erm* and HaloTag genes from the p100ptaHalo plasmid from the *Clj*-ptaHalo strain. Furthermore, the p100ptaHalo plasmid DNA has been integrated into the *Cac*-P4.5 genome, which was unexpected. There are relatively few reads in P4.5 that show integration, given the genome coverage of the sequencing data. Even in P1.5 and P2.5, where we see relatively high numbers of reads with plasmid DNA, we see a low number of reads indicating integration events in the *Cac* genome. The low read numbers are most likely due to the standard protocol employed for PacBio sequencing whereby reads higher than 6 kb were selected for sequencing. We speculate that free plasmid DNA survived the size selection process, whereas chromosomal DNA did not. As already argued, the coculture mediated cell fusion must have facilitated this successful transfer of plasmid DNA from *Clj*-ptaHalo to WT *Cac*. Second, the isolated PtP1.5 and PtP2.5 clones displayed a coculture-only phenotype of isopropanol and 2,3-butanediol production after five rounds of selection, designed to enrich for *Cac* cells. The possibility of a tight microcolony of distinct *Cac* and *Clj* cells can be discounted as the PtP1.5 strain could not survive the heat shock. Since these clones produce the characteristic solvents of *Cac* cells, the clones must contain the solventogenic genes, which are coded on the pSOL1 megaplasmid. Yet these clones are not heat-shock resistant. Survival and persistence of pure *Clj*-ptaHalo cells is unlikely due to the selection conditions (only glucose as a carbon substrate) during the five subcultures. The data therefore suggest that a large-scale exchange of chromosomal DNA has occurred between *Clj*-ptaHalo and the WT *Cac*, forming persistent & permanent hybrid cells. This allowed for the persistence of coculture phenotype in PtP1.5 and PtP2.5 clones.

**Figure 7.**
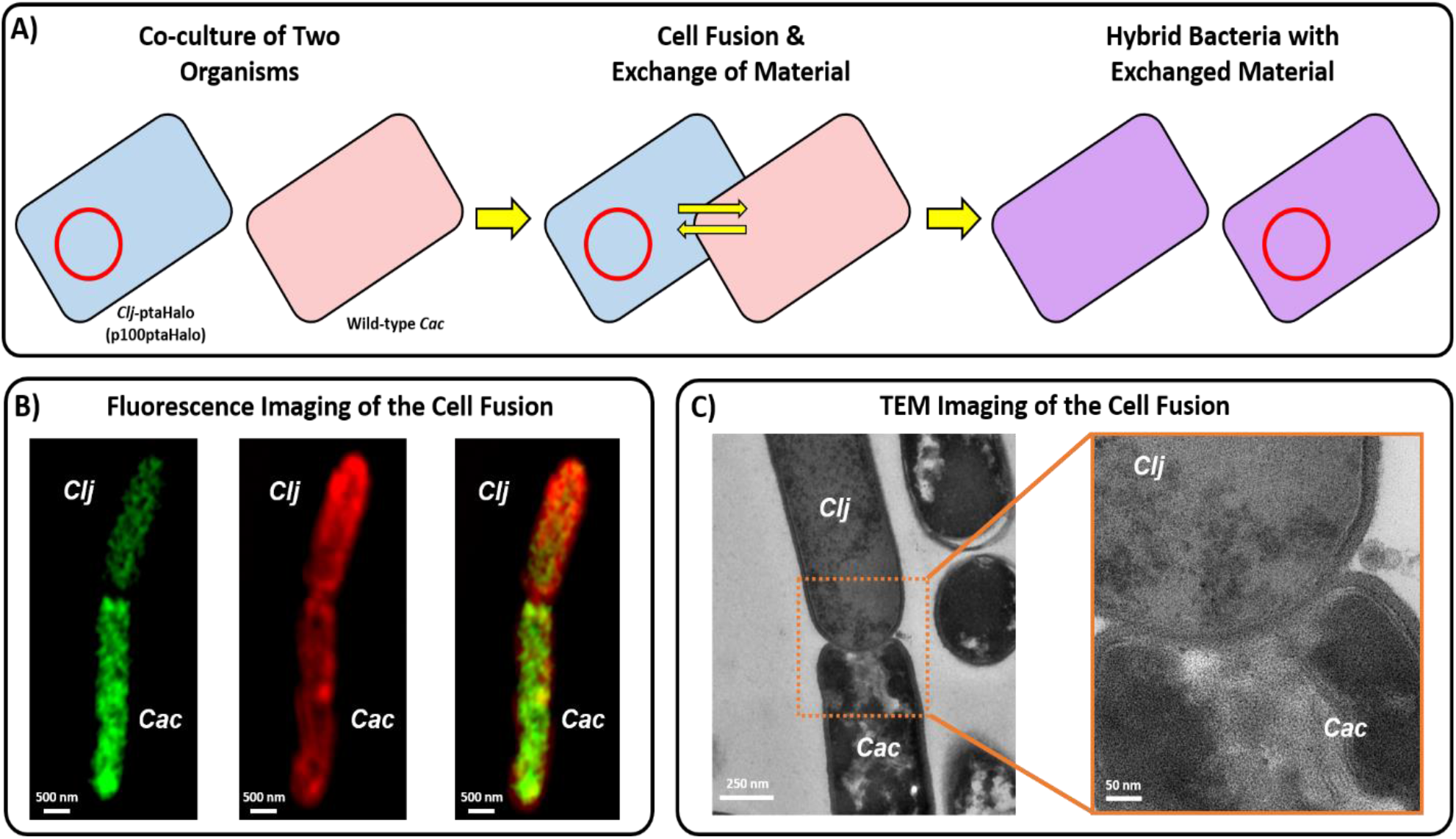
(A) Proposed mechanism of DNA transfer in the syntrophic coculture of *C. acetobutylicum* and *C. ljungdahlii*. Neither organism is known to possess any conjugation machinery. The restriction modification (RM) systems of the two organisms are incompatible, thus prevent any natural transformation from occurring. Thus, the observed DNA transfer must occur via the cell-to-cell fusion observed in the coculture system. (B) Fluorescence images (from Ref. ^1^) of heterologous cell-to-cell fusion between *C. acetobutylicum* and *C. ljungdahlii*, which facilitates the exchange of protein, RNA, and DNA between the two organisms. (C) Transmission-electron microscopy (TEM) images (from Ref. ^1^) of heterologous cell-to-cell fusion between the two organisms.

The exchange of plasmid, and possibly genomic, DNA in the coculture is strong evidence for a novel mechanism of HGT in bacteria. Control transformation experiments showed that the restriction-modification systems of *Cac* and *Clj* are incompatible (data not shown), making the DNA transfer through a transformation route not possible between these two organisms. There is also no evidence of transduction via prophage excision and transfer of DNA. *Clj* does have a large 51 kb prophage, and 3 more prophages scored as questionable or incomplete by the PHASTER online search tool (18, 19). *Cac* also has a 65 kb complete prophage and a 9 kb incomplete prophage, supported by previous Papoutsakis lab analysis (data not shown) (18, 19). If prophage excision was responsible for transfer of the plasmid DNA, we would expect that the plasmid first integrated into the *Clj* chromosome near a prophage region and then was excised and transferred to *Cac* along with prophage genes. PacBio analysis did not identify any reads with both plasmid sequence and *Clj* DNA in the P1.5 clone. Five reads containing both plasmid and *Clj* DNA were identified in the P2.5 clone; however, none of these reads mapped to regions of the *Clj* genome with prophage genes. No *Clj* DNA was detected in the P4.5 clone. Furthermore, there is no evidence of conjugative machinery in either *Cac* or *Clj*, meaning neither organisms can act as a donor cell during conjugation. Thus, the observed exchange of DNA in the coculture must have been facilitated through cell-to-cell fusion events that occur in the *Cac*-*Clj* cocultures. Cell fusion between these two organisms was found to facilitate large-scale exchange of protein and RNA (12), and, thus, it appears that plasmid-DNA exchange also took place during some fusion events. Formation of transient hybrid cells (exchange of protein and RNA only) may allow any transferred plasmid DNA to escape the restriction-modification of the recipient organism. At this point this mechanism is unclear. We did the best we could to examine the DNA content of isolated colonies originating presumably from single cells. To provide a definitive understanding of the events that took place and led to the observed phenotypes, one needs to be able to visualize the presence of chromosomal and plasmid DNA from the two parent cells in single cells during the selection process of Figure 1. This will require the development of new bioimaging technology not currently available for prokaryotic cells, such as the DNA PAINT technology (20-22).

Beyond the transfer of plasmid DNA, and possibly the generation of hybrid cells carrying, temporarily at least, two different chromosomes, heterologous fusion events would also facilitate the exchange of chromosomal DNA and notably mobile genetic elements, which are integrated in most prokaryotic genomes and can dynamically excise and reinsert (23, 24). For example, the *Cac* genome does contain an IS1595-like insertion sequence, which encodes a transposase (CA_RS07810). *C. acetobutylicum* ATCC 824 encodes 7 additional transposase genes (locus tags CA_RS01390, CA_RS03555, CA_RS03565, CA_RS03880, CA_RS08330, CA_RS13010, CA_RS18140). While all but one of the transposases either contain premature stop codons or encode only partial proteins, one or more of these transposases could be responsible for dynamic insertion and excision of the p100ptaHalo plasmid into the *Cac* genome. Evidence for movement of the plasmid or portions of the plasmid is seen in the PacBio data, as the plasmid is inserted in different locations in the genome in each generation of the P4 lineage. Thus, it is clear that heterologous fusion events can easily lead to novel cellular structures and phenotypes, such as those depicted in Figures 6 and S3, and such cellular entities may lead to novel evolutionary trajectories in prokaryotic biology, with important implications in environmental sciences and human and animal health.

Heteroresistance has been discussed and defined in the context of a single species, where “subpopulations of seemingly isogenic bacteria exhibit a range of susceptibilities to a particular antibiotic” (25). Additional qualifications have been proposed and the characteristic instability frequently associated with heteroresistance has been discussed (26). Here, all lineages of cultures shown in Figure 1 demonstrate a form of *Cac* heteroresistance to Erm, as *Cac* survives the presence of high concentrations (100 µg/mL) of Erm from the very first subculturing passage of all lineages. Moreover, we also observed that *Clj* cells displayed an unexpected phenotype in this syntrophic coculture: formation of colonies on the surface of 2xYTG plates, which, although not captured by the heteroresistance definition, involves acquisition of a distinct phenotypic trait not available to *Clj*. While the precise means by which these phenotypes occur remain to be elucidated, one could envision several plausible scenarios, all based on interspecies interactions. One possibility is the formation of tight microcolonies of *Clj*-ptaHalo with WT *Cac* cells, but this was discounted as discussed above. A second possibility would be immediate transfer of the p100ptaHalo plasmid to WT *Cac* cells, which while certainly possible, it appears less likely early on, such as during the PX.1 passages. Had that happened, subsequent, fast (24 hour) selective passages would have resulted in the isolation of *Cac* cells carrying the p100ptaHalo plasmid, while eliminating the *Clj*-ptaHalo cells, neither of which happened for lineages 1 and 2. A third possibility is the immediate formation of hybrid cells carrying both chromosomes and the p100ptaHalo plasmid, and there is evidence for such cells even in primary parent cultures (Figure 8 of Ref. (12)). Another possibility is the transient transfer of the Erm protein from the *Clj*-ptaHalo to WT *Cac* cells, as part of the massive exchange of proteins between these organisms under syntrophic co-culture conditions. While this is a most likely occurrence, such transient acquisition of the Erm protein would not have persisted in subsequent selective subcultures.

## MATERIALS AND METHODS

### Microorganisms and culture media

Monocultures of *C. acetobutylicum* (ATCC 824; *Cac*), the fluorescent *C. ljungdahlii* strain, *Clj*-ptaHalo (13), and the co-cultures thereof, were grown in Turbo CGM medium, as described previously (12, 14). Briefly, Turbo CGM used for *Clj*-ptaHalo mono-cultures was supplemented with 5 g/L fructose, and were grown in sealed bottles with 20 psig of H_2_/CO_2_ gas mixture (80/20%). Turbo CGM used for *Cac* mono-cultures, and co-cultures was supplemented with 5 g/L fructose and 80 g/L glucose, and were grown in unsealed glass bottles in the anaerobic chamber (12, 14).

### Monoculture preparation and growth

*Cac* frozen stocks were streaked onto 2xYTG plates and cultured in Turbo CGM to generate seed cultures (14). *Clj*-ptaHalo frozen stocks were inoculated into liquid Turbo CGM and passaged as needed to generate seed cultures, and were supplemented with erythromycin (100 μg/mL) to maintain the plasmid DNA. The culture pH was adjusted to 5.2 after 12 hours of growth with sterile deoxygenated 1 M NaOH to prevent acid death, as needed in *Cac* mono-cultures (14).

### Coculture setup

Co-cultures of *Cac* and *Clj*-ptaHalo were prepared as reported (12, 14). Briefly, 5 mL of exponentially-growing *Cac* seed cultures (OD_600_ of 1.0-2.0) were mixed with 90 mL of exponentially-growing *Clj*-ptaHalo seed cultures (OD_600_ of 0.4-0.6). The *Clj*-ptaHalo cells were spun down at 5,000 rpm and washed twice in the fresh Turbo CGM medium to remove any residual erythromycin, before using them for coculture setup. The cocultures used for the DNA transfer were prepared at the R of ∼10, where R is the ratio of *Clj* cells to *Cac* cells at the start of the co-culture (14), to ensure an excess of the plasmid-carrying *Clj*-ptaHalo cells (14). Co-cultures were performed in unpressurized static 100 mL glass bottles in the anaerobic chamber, with a total liquid volume of 30 mL. The pH of each coculture was adjusted to 5.2 with the sterile and deoxygenated NaOH after ∼12 hrs of growth to prevent the acid death (14). All coculture fermentations were performed using the Turbo CGM medium supplemented with 80 g/L of glucose and 5 g/L of fructose. No erythromycin was used in the starting parent cocultures.

### Selection procedure

After 24 hrs of the initial (parent) coculture, samples were collected for selection in order to isolate any *Cac* cells that acquired the p100ptaHalo plasmid (carrying the HaloTag gene and the erythromycin resistance gene; see ref. (13) for details) from the *Clj*-ptaHalo. The selection was done in liquid medium and solid plates. The liquid selection medium was the Turbo CGM medium containing 80 g/L of glucose, no fructose and 100 μg/mL of erythromycin. The liquid selection cultures were done in unsealed 100 mL bottles in the anaerobic chamber. The plate selection was done on 2xYTG plates, containing 5 g/L of glucose, no fructose, and 100 μg/mL of erythromycin. During the liquid and solid selection, the presence of only high glucose concentration (a *Cac* substrate), but no fructose (*Clj* substrate), was expected to enrich the original cocultures samples in *Cac* cells, while eliminating *Clj*-ptaHalo over the course of the selection. The selection media also contained the erythromycin (100 μg/mL) in order to eliminate WT *Cac* cells during the selection process, and over time isolate only *Cac* cells that acquired the plasmid p100ptaHalo DNA (or parts of it) in the coculture. The subcultured clones isolated at any point during the selection process were designated as PX.#, where ‘X’ represents the different starting (parent) co-cultures, while the ‘#’ represents the specific subculture stage (passage). Plate subculture selection is indicated by PtPX.#. To start the selection process, 5 mL samples from each mother coculture were washed in Turbo CGM medium (80 g/L of glucose only, no fructose) and transferred to 25 mL of the liquid selection medium. The coculture samples were washed to remove any fructose left over from the coculture growth medium. This was the 1^st^ selection passage P*X*.1 (Figure 1). After 24 hrs of growth, 5 mL samples from each P*X*.1 culture were collected, washed, and transferred to fresh 25 mL of the selection liquid medium (passage P*X*.2). After 24 hrs of incubation, samples from P*X*.2 cultures were steaked onto 2xYTG selection plates (PtP*X*.3) to begin isolating and testing single colonies of each clone. The selection plates developed colonies after 2 days of incubation at 37°C, except the selection plate PtP3.3. Multiple colonies (8-10) were picked from each selection plate, and were cultured in the liquid selection medium (80 g/L glucose, 100 μg/mL erythromycin). Half of the selected colonies were heat-shocked at 80°C for 10 minutes (per standard *Cac* culture techniques) to check whether the colonies from each plate were able to sporulate. All colonies that grew in the liquid selection medium were streaked again on the 2xYTG selection plate (plate PtP*X*.4), and the process was repeated one more time (plate PtP*X*.5) to further enrich each clone. Colonies from plates PtP1.5, PtP2.5, and PtP4.5 were grown in the selection liquid medium to generate cells for microscopy, flow-cytometry, HPLC, and PCR analysis.

### Transmission Electron Microscopy (TEM)

Samples from cultures PtP1.5, PtP2.5, and PtP4.5 were collected after 24 hrs of growth, fixed in 2% glutaraldehyde and 2% paraformaldehyde in 0.1M sodium cacodylate buffer (pH 7.4) and stored at 4°C until further processing. The TEM sample processing and imaging was performed as described previously (12).

### Confocal Fluorescence Microscopy

Samples from the PtP4.3 cultures were collected after 24 hrs of growth, and labeled with the HaloTag-specific Janelia Fluor®646 red ligand as described previously (12, 13). Labeled cells were placed in Nunc Lab-Tek chamber slides coated with poly-L-lysine. Cells were incubated for 1 hr to immobilize cells on the poly-L-lysine coating. After 1 hr, the chamber was rinsed with PBS thrice to remove excess cells, as described previously (12, 13). Immobilized cells were imaged using Elyra PS.1 super-resolution microscope (Carl Zeiss). Each sample was imaged using a 63x/1.4 oil objective, as described previously (12, 13).

### Flow cytometry and fluorescent labeling of cells

Samples from passage cultures P1.5, P2.5, and P4.5 were collected, and labeled with the HaloTag-specific Janelia Fluor®646 red ligand as described previously (12, 13) to determine if the isolated cells still produced the HaloTag protein. The flow cytometry analysis was performed as described previously (12, 13).

### HPLC metabolite analysis

Colonies from isolated strains P1.5, and P4.5 were grown in the liquid selection medium (Turbo CGM, 80 g/L glucose, no fructose, 100 μg/mL erythromycin) for 40 hrs, with pH control at 12 hrs, to determine the fermentation profile of each isolated passage line. The samples were collected approximately every 10-12 hrs for the HPLC analysis. The HPLC analysis was performed as described previously (14).

### Plasmid isolation from isolated clones

Cell samples of wild-type *Cac*, wild-type *Clj, Cac*-ptaHalo, *Clj-*ptaHalo, and clones PtP1.5, PtP2.5, and PtP4.5 were used for plasmid isolation, using the NucleoSpin Plasmid Mini Kit (Macherey-Nagel) according to manufacturer’s protocol. To test for the presence of a complete p100ptaHalo plasmid, the resulting plasmid preparations from each clone were transformed into chemically-competent NEB 5-alpha *E. coli* cells per the standard transformation protocol. Following transformation, E. coli cells were incubated at 37°C for one hour, after which 150 μL of each transformation were plated on LB plates supplemented with 100 μg of ampicillin (p100ptaHalo plasmid contained Amp^R^ marker for *E. coli* transformation). The LB plates were incubated at 37°C for 24 hrs to allow *E. coli* colonies to develop. Plasmid preparations from clones PtP1.5 and PtP2.5 produced *E. coli* colonies, indicating that the complete plasmid DNA was present in those samples. Plasmid preparations from PtP4.5 did not produce any colonies, indicating the complete plasmid was not present in this clone.

### PCR analysis of genomic and plasmid DNA

Genomic and plasmid DNA was extracted from isolated strains P1.5, and P4.5, as well as WT *Cac* (negative control), WT *Clj* (negative control) and *Cac*-ptaHalo (positive control) for PCR analysis. Genomic DNA was extracted using the DNeasy Blood & Tissue Kit (Qiagen, Germany), following the procedure for Gram positive bacteria (14). Plasmid DNA was extracted using the NucleoSpin Plasmid DNA Kit (Macherey-Nagel, Germany).

Genomic samples extracted from isolated strains P1.5 and P4.5 were first tested by PCR using primers to target five selected *Cac* genes *(adc, ctfa, ctfb, adhe, thl*) and three *Clj* genes (*sadh, 23bdh, rho*) to determine if pure *Cac* was isolated in each line during the selection process. Primers used to screen for the selected *Cac* and *Clj* genes have been used previously (14). PCR was performed using the green 2X Taq polymerase master mix (Fisher, MA). Each reaction was performed under the following conditions: initial 5 min denaturation at 95°C; followed by 25 cycles of 30 sec denaturation at 95°C, 30 sec annealing at 65°C, and 30 sec extension at 72°C; finished with 5 min extension at 72°C. All primer sets were designed to have the same annealing temperature of 65°C.

Genomic and plasmid DNA samples from isolated strains P1.5 and P4.5 were also tested for the presence of the HaloTag, and erythromycin resistance (*erm*) genes. Three primer sets were designed for each gene, where the left pair (L) spanned the 5’-end of the gene and the plasmid backbone DNA located to the left of the gene, the middle pair (M) spanned a region of each gene in the middle, and the right pair (R) spanned the 3’-end of the gene and the plasmid backbone DNA located to the right. This is summarized visually in the Figure 4, while the primer sequences are shown in Table S2. PCR was performed using the green 2X Taq polymerase master mix (Fisher, MA). Each reaction for the *erm* gene (L, M, R) was performed under the following conditions: initial 5 min denaturation at 95°C; followed by 25 cycles of 30 sec denaturation at 95°C, 30 sec annealing at 53°C, and 35 sec extension at 72°C; finished with 5 min extension at 72°C. Each reaction for the HaloTag gene (L, M, R) was performed under the same conditions as above, except the annealing temperature was 53°C for these primer pairs (see Table S2).

### Whole-genome PacBio Sequencing and Bioinformatics

High molecular weight (HMW) DNA was isolated from Cac-P4.5 cells (originating from the PtP4.5 selection plate) using the MagAttract HMW DNA Kit (Qiagen) according to the manufacturer’s instructions. Single molecule real time sequencing was performed at the University of Delaware DNA Sequencing and Genotyping Center. DNA libraries were constructed according to the PacBio standard protocol. SMRTbell DNA libraries were constructed as described previously (27). Briefly, DNA libraries were constructed according to the PacBio standard protocol. Both libraries were size-selected starting at 6 kb and with an average library size of 10 kb, as measured by Fragment Analyzer (Advanced Analytical Technologies, Inc.). DNA Sequencing was performed on PacBio Sequel II Single-Molecule Sequencer (Pacific Biosciences, Menlo Park, CA) instrument using P4-C2 chemistry, mag-bead loading and 3-h movie time. Reads from each clone were assembled into contigs using SMRT Analysis version 10.1 through the SMRT Portal. The CCS (Circular Consensus Sequence) tool from PacBio SMRT Tools v10.1 was used to calculate the consensus sequences from the subreads in the PacBio data for each sample. The CCS reads were then aligned to the *erm* gene, HaloTag gene, and the entire p100ptaHalo plasmid using BLASTn (v2.11.0) (28). Reads aligning to more than 500 contiguous base pairs of either the *erm* gene, HaloTag gene, or the plasmid were then mapped to the *Cac* reference genome (GCF_000008765.1) to identify possible integration sites using minimap2 (v2.1) (29). These reads were also aligned to the *Cac* reference genome using BLASTn to filter by alignment length and percent identity. In addition, the presence of *Clj* DNA in each sample was examined by mapping the reads to the *Clj* reference genome (GCF_000143685.1) and identifying which reads mapped to *Clj* better than *Cac* and to *Clj* only. The presence of *Clj* DNA with plasmid was examined by aligning the PacBio reads that contained more than 500 contiguous base pairs of plasmid to the *Clj* reference genome (GCF_000143685.1) using BLASTn. Online tools ICEFinder, Mobile Element Finder, and PHASTER were used to search the reference genomes of *C. acetobutylicum* ATCC 824 and *C. ljungdahlii* DSM 13528 genomes for integrative and conjugative elements, mobile genetic elements and prophages, respectively (16-19).

## Acknowledgements

This work was supported by a grant from the Army Research Office (award W911NF-19-1-0274) and unrestricted institutional funds available to ETP. Microscopy equipment was acquired with a shared instrumentation grant (grant S10 OD016361), and access was supported by the NIH-NIGMS (grant P20 GM103446), by the NSF (grant IIA-1301765) and by the State of Delaware. Support from the University of Delaware CBCB Bioinformatics Core Facility and use of the BIOMIX compute cluster was made possible through funding from Delaware INBRE (NIH NIGMS P20 GM103446), the State of Delaware, and the Delaware Biotechnology Institute. We thank Shannon Modla & Jeffrey L. Caplan of UD Bio-Imaging Center for assistance in TEM and confocal microscopies, Olga Shevchenko of UD DNA Sequencing & Genotyping Center for assistance with PacBio sequencing, and Madolyn Macdonald & Shawn Polson of UD Center for Bioinformatics and Computational Biology for assistance with PacBio sequencing data analysis.

## AUTHOR CONTRIBUTIONS

ETP conceived the project. ETP and KC designed the experiments; KC with ETP performed culture experiments. GJG performed DNA isolation and data analysis for PacBio sequencing. ETP, KC, and GJG analyzed the data and wrote the manuscript.

