## Supplemental Information for "Novel mechanism of plasmid-DNA transfer mediated by heterologous cell fusion in syntrophic coculture of *Clostridium* organisms"

**Table S1.** Phenotypic checklist of the parent strains used for the starting coculture (*Clj*-ptaHalo and WT *Cac*), and the observed phenotype in PtP1.5 and PtP2.5 subculture clones, which show evidence of genomic DNA transfer from *Clj*-ptaHalo to WT *Cac*.

| ***Characteristic*** | ***Clj*-ptaHalo** | **Wild-type *Cac*** | **Clones from PtP1.5 Plate** | **Clones from PtP2.5 Plate** | |
|---|---|---|---|---|---|
| *Heat-shock survival* | **No** | **Yes** | **No** | **No** | ***Cac* Specific** |
| *Growth on glucose only* | **No** | **Yes** | **Yes** | **Yes** | |
| *Growth on 2xYTG plate surface* | **No** | **Yes** | **Yes** | **Yes** | |
| *Prod. of butanol, acetone* | **No** | **Yes** | **Yes** | **Yes** | |
| *Growth on erythromycin* | **Yes** | **No** | **Yes** | **Yes** | **Plasmid Specific** |
| *HaloTag fluorescence* | **Yes** | **No** | **Yes** | **Yes** | |
| *Production of isopropanol* | **No** | **No** | **Yes** | **Yes** | **Co-culture Specific** |

**Table S2** Primer pairs used for the HaloTag and the erythromycin resistance ($Erm^R$) gene assay in the isolated strains *Cac*-P1.5 and *Cac*-P4.5. Primers were designed using Primer3Plus online tool (https://www.bioinformatics.nl/cgi-bin/primer3plus/primer3plus.cgi). The annealing temperature ($T_A$), the expected product size, the directionality, and the sequence of each primer are listed.

| **Gene** | **Reaction** | **Size (bp)** | **For/Rev** | **Primer Sequence (5' – 3')** |
|---|---|---|---|---|
| *erm* Gene ($T_A$=50°C) | Left (L) | 345 | F | AGATACTGCACCCCCTGAAC |
| | | | R | TTTCGTTATGAAATGGGTTAACAA |
| | Middle (M) | 328 | F | TAATGCCAATGAGCGTTTTG |
| | | | R | TGAAATCGGCTCAGGAAAAG |
| | Right (R) | 480 | F | CTTTTCCTGAGCCGATTTCA |
| | | | R | TATTTCACTTAGGCATTTCACG |

| | | | | |
|---|---|---|---|---|
| HaloTag Gene ($T_A$=53°C) | Left (L) | 470 | F | GGTCGTAGAGCACACGGTTT |
| | | | R | CGTAATGAGGGTCAAATGGG |
| | Middle (M) | 344 | F | CCCATTTGACCCTCATTACG |
| | | | R | CTCTTTCAGGGTTTCTCTTTGC |
| | Right (R) | 548 | F | TTCTGAGATTGCAAGGTGGTT |
| | | | R | TTTTTGTGATGCTCGTCAGG |

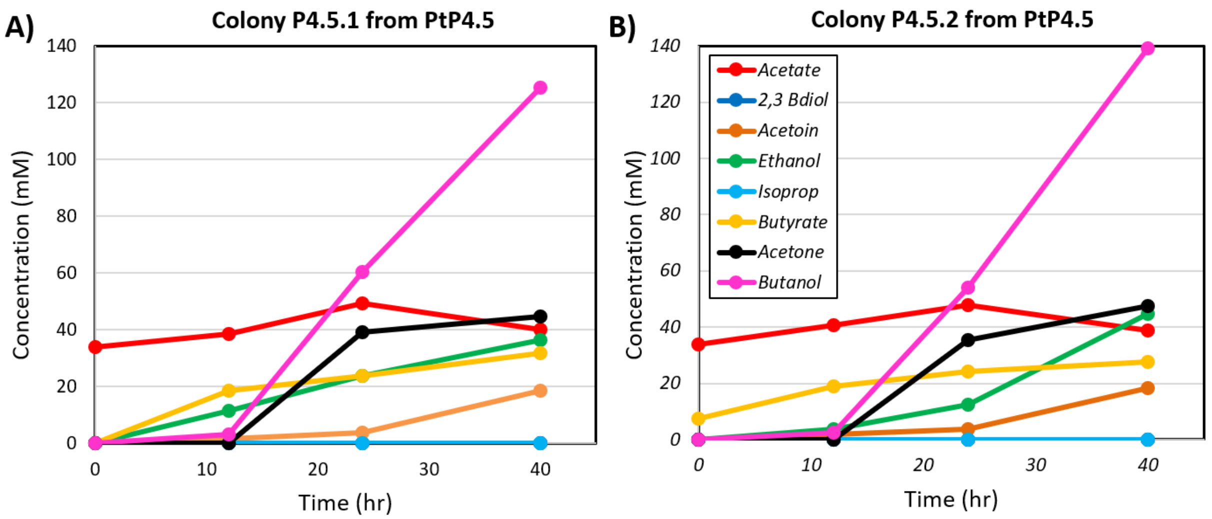


**Figure S1.** Metabolite profile of the clones isolated from subculture selection plate PtP4.5. Panels (A) and (B) show results from two colonies picked from the plate PtP4.5 and subcultured in selective liquid medium (glucose, Erm, no fructose). These clones exhibited the pure *Cac* phenotype of butanol, acetone, and acetoin production. No isopropanol was detected, indicating the selection process removed all *Clj*-ptaHalo cells.

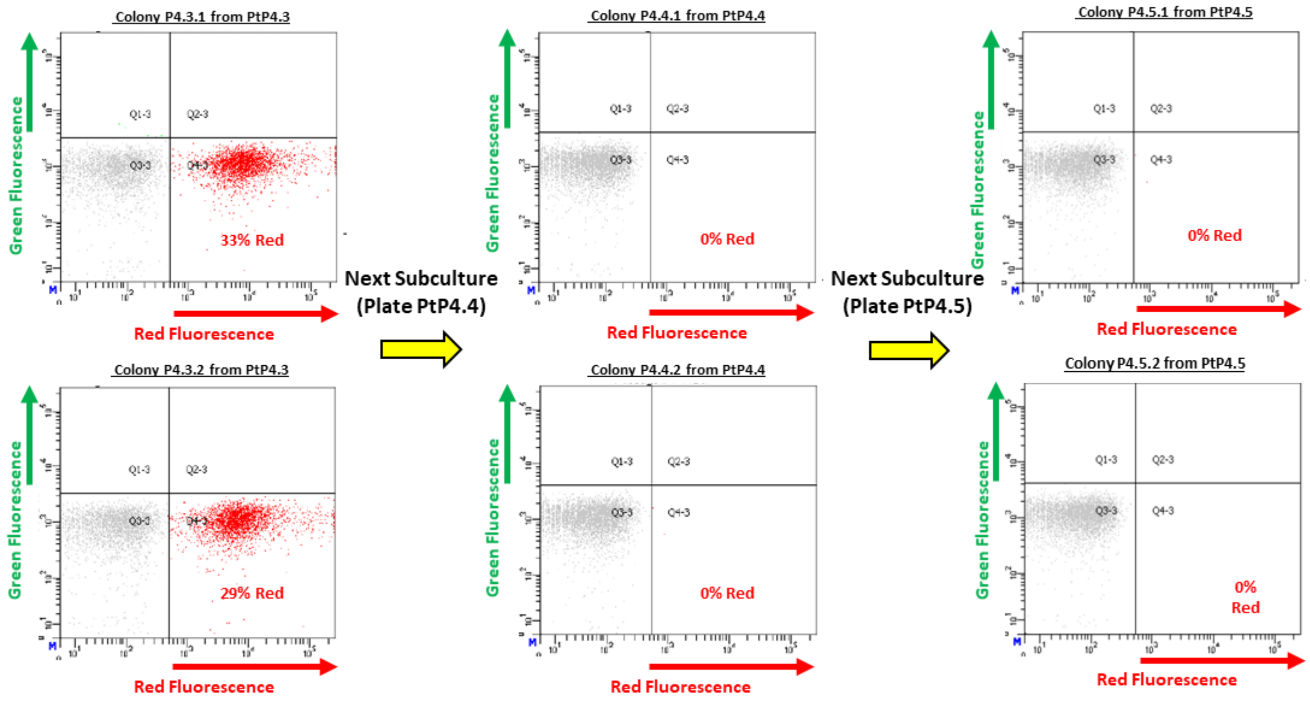


**Figure S2.** Flow cytometric analysis of clones isolated from subculture selection plates PtP4.3, PtP4.4, and PtP4.5. Two individual colonies were picked from each selection plate (PtP4.3, PtP4.4, and PtP4.5), grown in the selective medium (glucose, Erm, no fructose) and analyzed. Cells were labeled with the red Janelia Fluor®646 ligand (see Methods). PtP4.3 cells exhibited red fluorescence in approximately 30% of the population. Starting with the plate selection PtP4.4, no red fluorescent cells were detected using flow cytometry, indicating the lack of – detectable by flow cytometry – expression of the HaloTag protein in PtP4.4 and PtP4.5 cells.

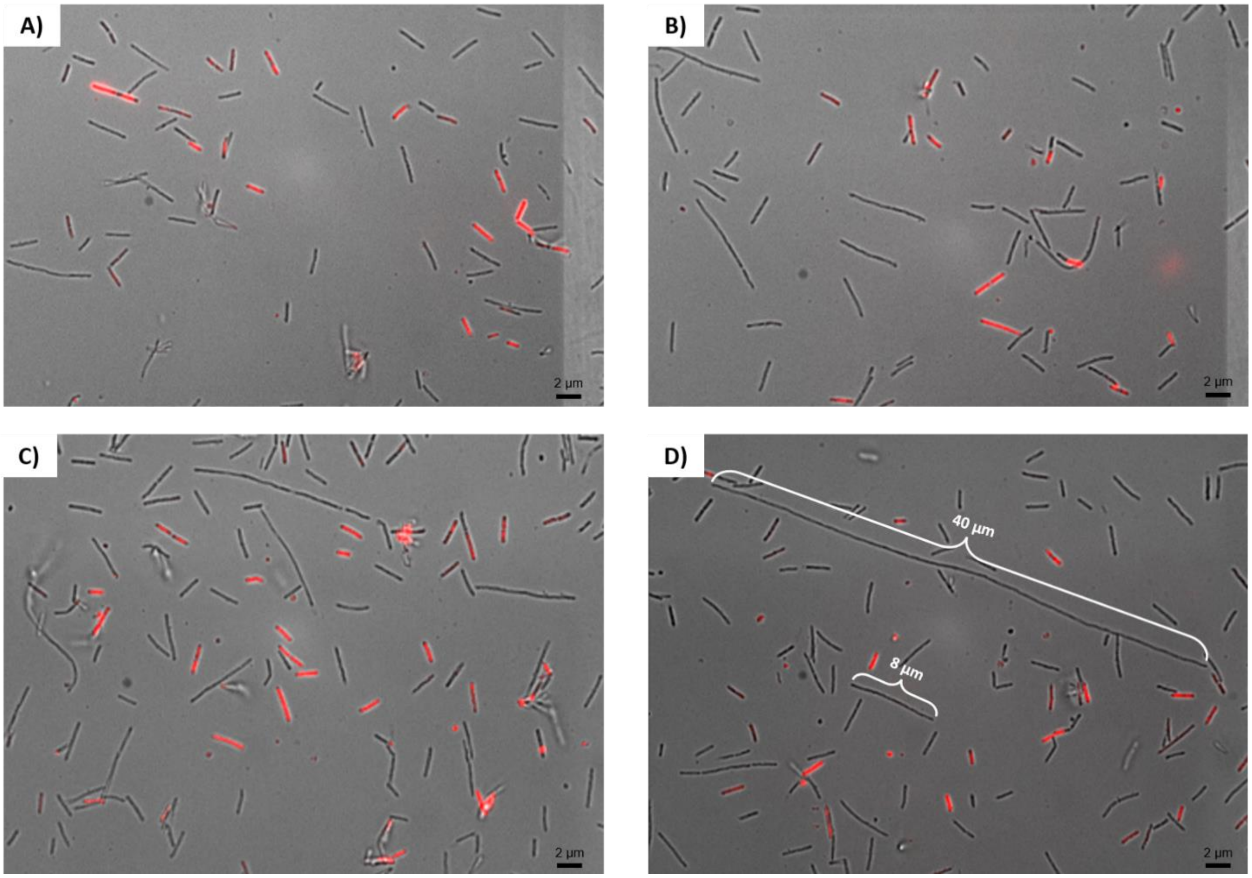


**Figure S3.** Confocal fluorescence imaging of the PtP4.3 cells. Cells were labeled with the red Janelia Fluor®646 ligand (see Methods). Approximately a third (33%) of the PtP4.3 cells showed red fluorescence, similar to what was assessed by flow cytometry analysis (Figure S2), indicating the presence of the functional HaloTag protein. Only the smaller, normal-size cells show red fluorescence. The abnormal long cells do not express HaloTag protein sufficient for detection of red fluorescence or they express none. All panels show an enrichment for abnormally long cells (> 4-6 μm) in excess of normal length of about 2-3 μm of wild-type *Cac* or *Clj* cells. For example, panel (D) shows elongated cells ranging from 8 μm up to 40 μm in length, none with red fluorescence.

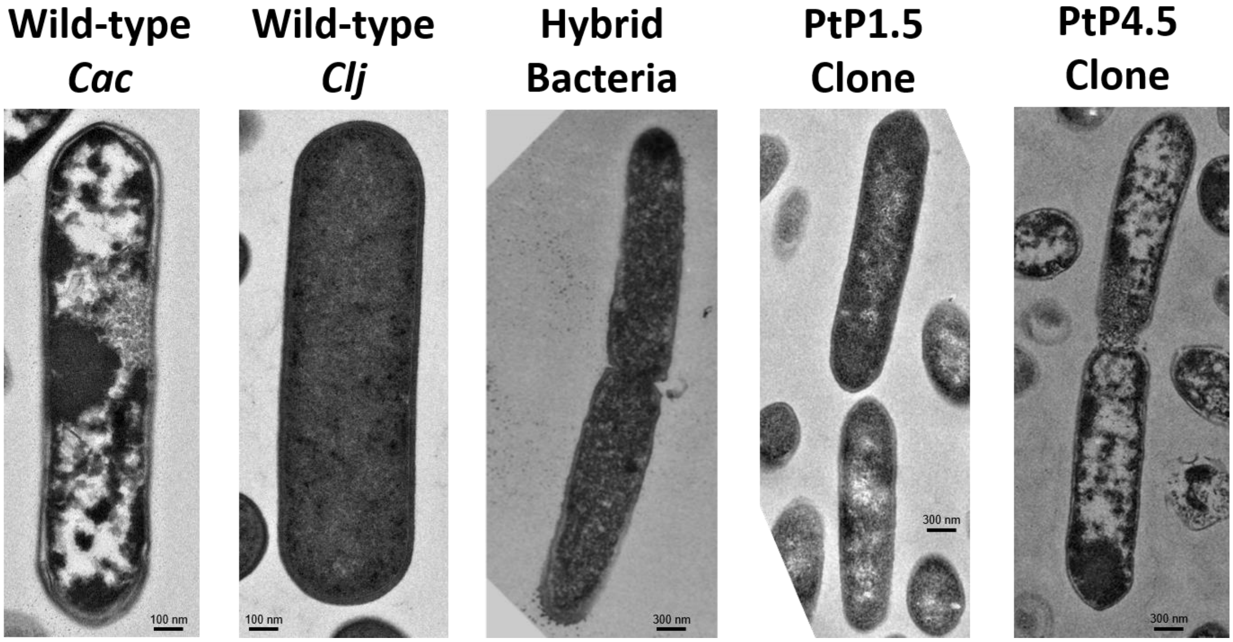


**Figure S4.** Comparison of morphologies of various strains and species. All cells were examined using TEM after 24 hrs of growth. Wild-type *C. acetobutylicum* (*Cac*) exhibited large translucent regions within its cytoplasm called granulose, which form as part of its sporulation program. *C. ljungdahlii* (*Clj*) cells remain homogenously electron dark, as is typical of vegetative cells with no signs of any differentiation/sporulation. The morphology of hybrid bacterial cells, observed in cocultures between *Cac* and *Clj*, showed a mixed morphology that appeared to be a mixture of homogenous *Clj* and differentiated *Cac* morphologies (1). TEM imaging of the PtP1.5 clone cells showed a mixed morphology unlike either *Cac*- or *Clj*-cell morphologies. This is strong evidence that the PtP1.5 clone cells were hybrid bacterial cells containing chromosomal DNA of the two parent organisms. TEM imaging of the PtP4.5 clone cells similar to those of the wild-type *Cac* cells. This is strong evidence that the PtP4.5 cells are wild-type *Cac* cells that successfully acquired the plasmid DNA from *Clj*-ptaHalo cells during the co-culture.

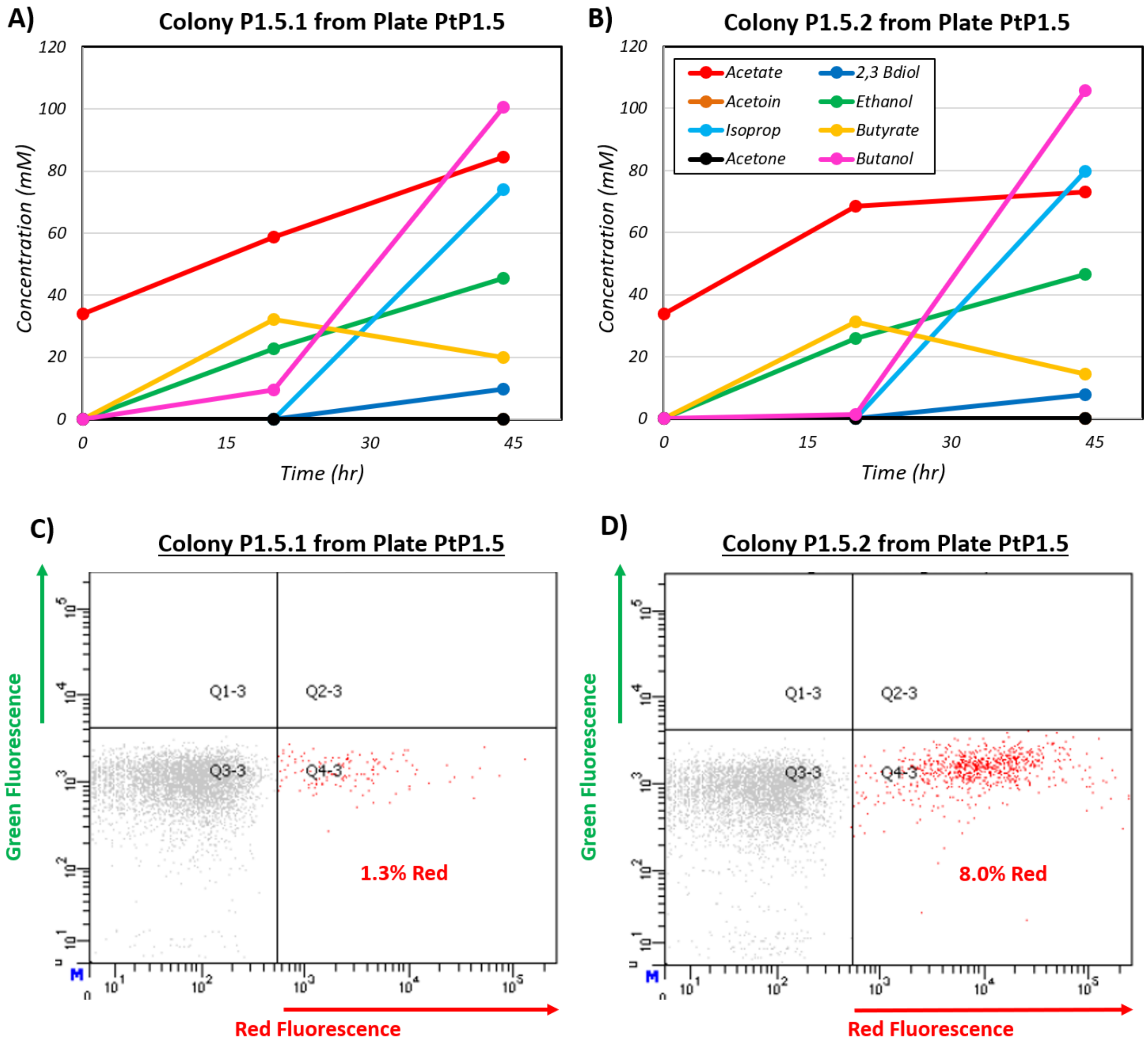


**Figure S5.** (A) and (B): Metabolite profile of two PtP1.5 clones. Two individual colonies from plate PtP1.5 were picked and cultured in the liquid selective medium. Two resulting subcultures were used for metabolite and fluorescence analysis. (C) and (D): Flow cytometric analysis of PtP1.5 clones. Cells were labeled with the red Janelia Fluor®646 ligand (see Methods).
